# LncMamba: A Deep Learning Model for lncRNA Localization Prediction Based on the Mamba Model

**DOI:** 10.1101/2025.03.29.646080

**Authors:** Baixiang Huang, Yu Luo, Yumeng Zhuang, Songyan He, Chunxin Yuan

## Abstract

Accurate prediction of long non-coding RNA (lncRNA) subcellular localization is crucial for understanding its biological functions. In this study, we propose a novel deep learning framework, LncMamba, which utilizes a two-layer FPN network for multi-scale feature extraction and introduces the Mamba network to lncRNA localization prediction tasks for the first time. Based on this, we improved the localization-specific attention mechanism, allowing the model to more effectively focus on key sequence motifs related to localization. Additionally, through statistical analysis of localization motifs, we revealed the potential relationship between nucleotide motifs and lncRNA subcellular localization.The code is available at: https://anonymous.4open.science/r/lncMamba-731F.

## Introduction

Long non-coding RNAs (lncRNAs) are RNA transcripts longer than 200 nucleotides that do not code for proteins. The subcellular localization of lncRNAs is closely related to their biological functions. lncRNAs located in the nucleus may regulate gene expression, while those distributed in the cytoplasm are often associated with protein translation or signaling pathways [24]. For instance, in the cytoplasm, lncRNAs can interact with mRNA to regulate the translation process [6] lncRNAs are also involved in various stages of cancer development. These lncRNAs interact with DNA, RNA, protein molecules, and/or their combinations, serving as essential regulators of chromatin organization, transcription, and post-transcriptional regulation. Their dysregulation endows cancer cells with the ability to initiate, grow, and metastasize [21]. Therefore, the correct localization of these lncrnas plays a crucial role in the development of targeted drugs and the early treatment of cancer. The ability of lncRNAs to interact with various molecular species highlights the importance of understanding lncRNA localization as a key to predicting its function [4].

Traditionally, experimental methods such as RNA fluorescence in situ hybridization (FISH) have been used to determine the localization of lncRNAs. However, these techniques are labor-intensive and unsuitable for high-throughput analysis [17]. Therefore, developing accurate and efficient computational methods for predicting lncRNA subcellular localization is crucial for biological research.

Currently, many research methods have been widely applied in the field of lncRNA localization prediction. From traditional machine learning methods, for example, LncSL [31] integrates multiple feature selection algorithms, uses stacking strategies, and automatically selects appropriate machine learning models. Locate-R [15] uses n-gap oligomer compositions and oligomer compositions as features, employing local deep support vector machines. lncLocPred [29] utilizes sequence features such as k-mers, trigrams, and PseDNC to develop a logistic regression-based machine learning predictor. LncPred-IEL and LncPred-ANEL [20] are lncRNA prediction methods based on feature ensemble learning strategies, and they perform well in cross-species lncRNA prediction. lncLocator [30] constructs classifiers by feeding k-mer features and high-level abstract features generated by unsupervised deep models into support vector machines (SVM) and random forests (RF).

With the rise of deep learning in recent years, more studies have started using deep neural networks for lncRNA localization prediction. For instance, EL-RMLocNet [1] optimizes prediction performance and interpretability using the GeneticSeq2Vec statistical representation learning scheme and attention mechanisms, proposing an LSTM-based model. LncLocFormer [23] employs eight Transformer blocks to capture long-range dependencies within lncRNA sequences and utilizes a subcellular localization-specific attention mechanism. The CFPLncLoc model [18] predicts multi-label lncRNA subcellular localization by using chaotic game representation (CGR) images of lncRNA sequences and a CNN classifier.

DeepLncLoc [26] encodes complete lncRNA sequences using subsequence embeddings and applies a text convolutional neural network (textCNN) to learn high-level features and perform prediction tasks. GraphLncLoc [11] converts lncRNA sequences into De Bruijn graphs and uses graph neural networks for feature extraction, retaining the local sequence order information. SGCL-LncLoc [10] builds upon converting lncRNA sequences into graphs and introduces supervised contrastive learning and global attention pooling mechanisms to enhance discriminability and interpretability in lncRNA subcellular localization.

In this study, we propose a novel deep learning model called lncMamba for multi-label lncRNA subcellular localization prediction. lncMamba integrates a dual-layer FPN module [12] that efficiently captures multi-scale features and leverages the Mamba network’s strength in handling long sequences—ensuring that both local sequence details and global context are effectively represented. Based on the LncLocFormer [23] framework and results from LncLocator 2.0 [14], we improved the subcellular localization attention mechanism, enabling the model to more effectively focus on important sequence motifs related to localization. We conducted extensive experiments to evaluate the effectiveness of lncMamba. We compared it with various machine learning and deep learning baseline models as well as existing predictors. The results clearly show that lncMamba outperforms these models. Additionally, we performed ablation studies to analyze the impact of different components on prediction performance, confirming the contribution of each component. We also performed statistical analysis to explore the relationship between nucleotide motifs and lncRNA subcellular localization. The flowchart of our method is shown in Figure 1. We first directly obtained two publicly available datasets, performed data preprocessing, then input the data into our deep learning network, and finally conducted important motif analysis.

**Fig. 1.**
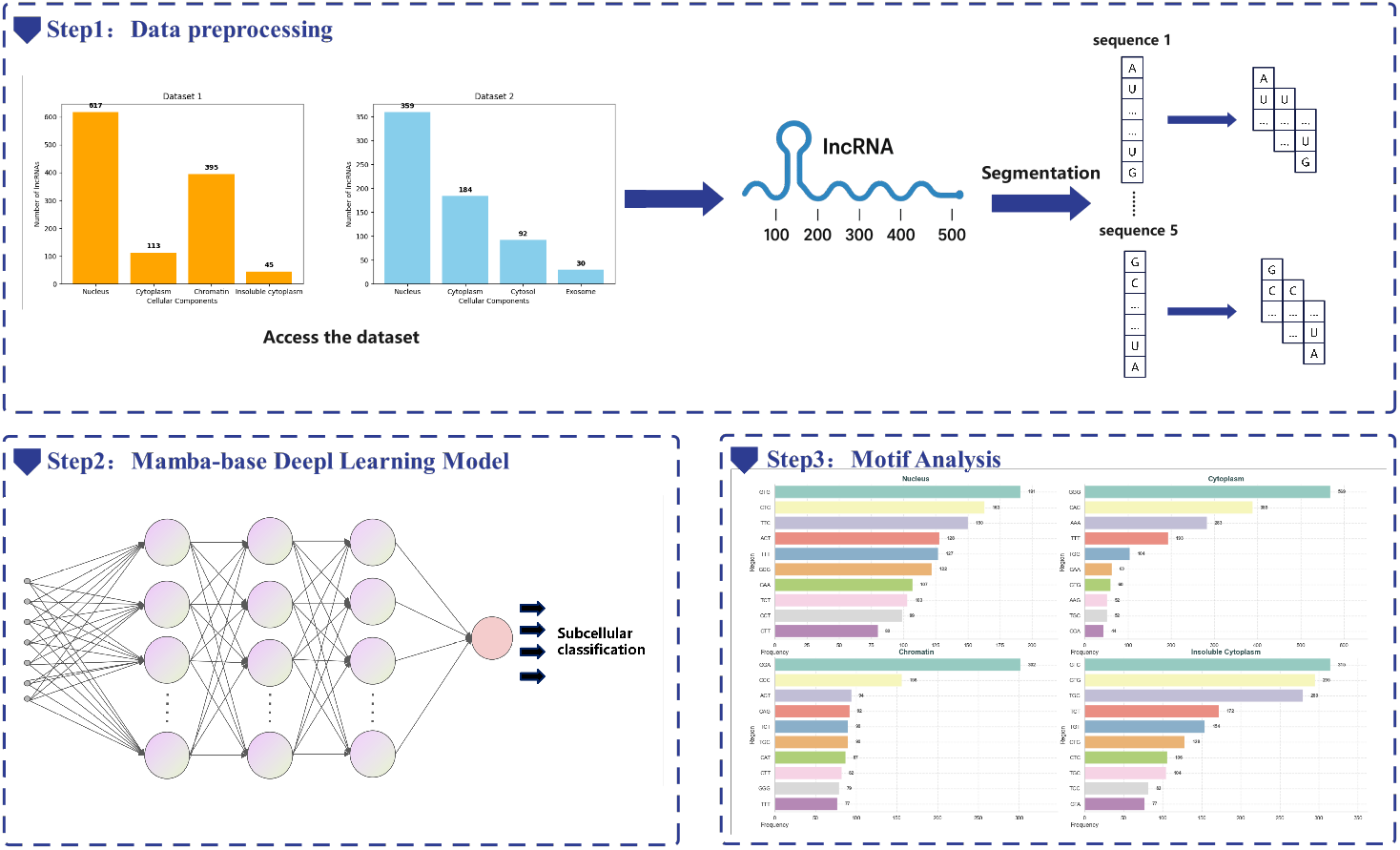
Flowchart of our method.

## Materials and methods

### Dataset

We directly utilized the benchmark datasets constructed by Zeng et al. [28] (LncLocFormer) and Bai et al. [2] (ncRNALocate-EL), designating them as Dataset 1 and Dataset 2 respectively. Dataset 1 contains four subcellular categories: Nucleus, Cytoplasm, Chromatin, and Insoluble Cytoplasm. Dataset 2 comprises four categories: Nucleus, Cytoplasm, Cytosol, and Exosome. We statistically analyzed the sample count (*N*) and average sequence length (*µ*) for each subcellular category, as shown in Fig. 2. All samples in both datasets originate from Homo sapiens. Following a 9:1 split ratio, the 729 samples of Dataset 1 were allocated to the training set, with the remaining 82 samples serving as an independent test set.As shown in Fig. 2, the majority of lncRNA sequences in the datasets are in the order of 10^3^ in length. Furthermore, the sequence lengths in Dataset 2 are generally longer compared to those in Dataset 1.

**Fig. 2.**
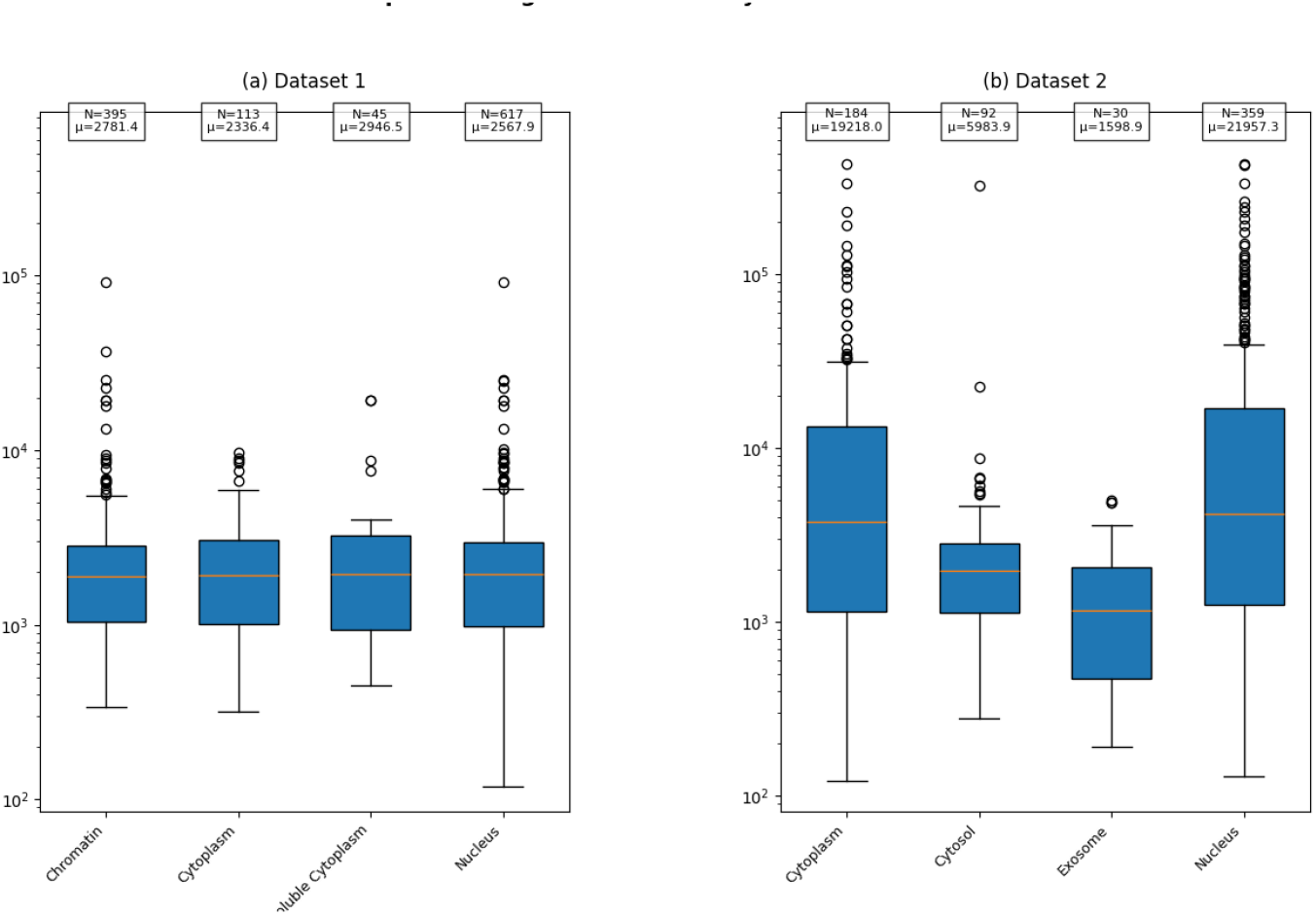
Statistical characteristics of benchmark datasets.

### LncMamba Architecture

#### Preliminaries: State Space Mode

The state space model is derived from control theory, specifically linear time-invariant (LTI) systems, as described in the foundational work by Kalman [9]. It maps a multidimensional input signal *X*(*t*) to an *N* -dimensional latent state *h*(*t*), which is then projected into a scalar output signal *y*(*t*). This process can be described by the following system of ordinary differential equations (ODEs)[16]):

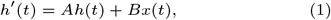

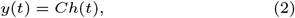

where *A* ∈ ℝ^*N ×N*^ is the state transition matrix, *B* ∈ ℝ^*N*^ is the input matrix, and *C* ∈ ℝ^*N*^ is the projection matrix.

To adapt this model to discrete inputs, commonly encountered in deep learning applications such as text sequences, we discretize the continuous system using the zero-order hold (ZOH) method [5]. This process involves introducing a learnable time scale parameter Δ, transforming the continuous state space model (SSM) into a discrete version. The discretization process is as follows:

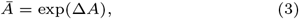

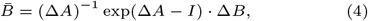

After discretization, the equations become:

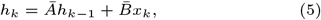

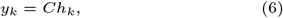

where *Ā* and 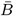 are the discretized versions of the state transition matrix *A* and input matrix *B*, respectively. *h*_*k*−1_ refers to the state at the previous time step, while *h*_*k*_ represents the current state.

#### Architecture Overview

The architecture of the LncMamba model is shown in Figure 3. It consists mainly of an embedding layer, a two-layer FPN module, 1 Mamba2 block, an improved localization-specific attention mechanism, and a classifier.

**Fig. 3.**
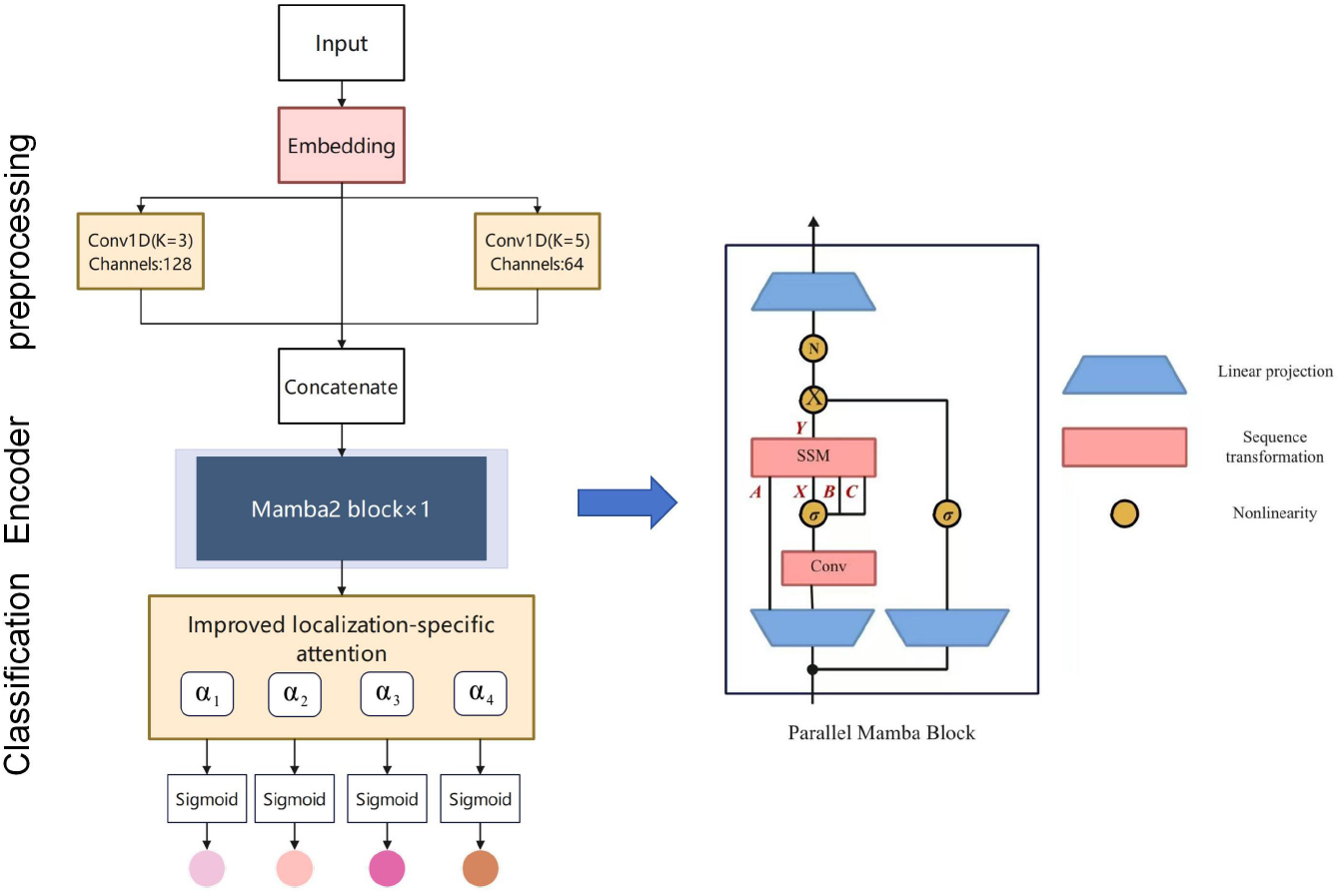
LncMamba model architecture

**Fig. 4.**
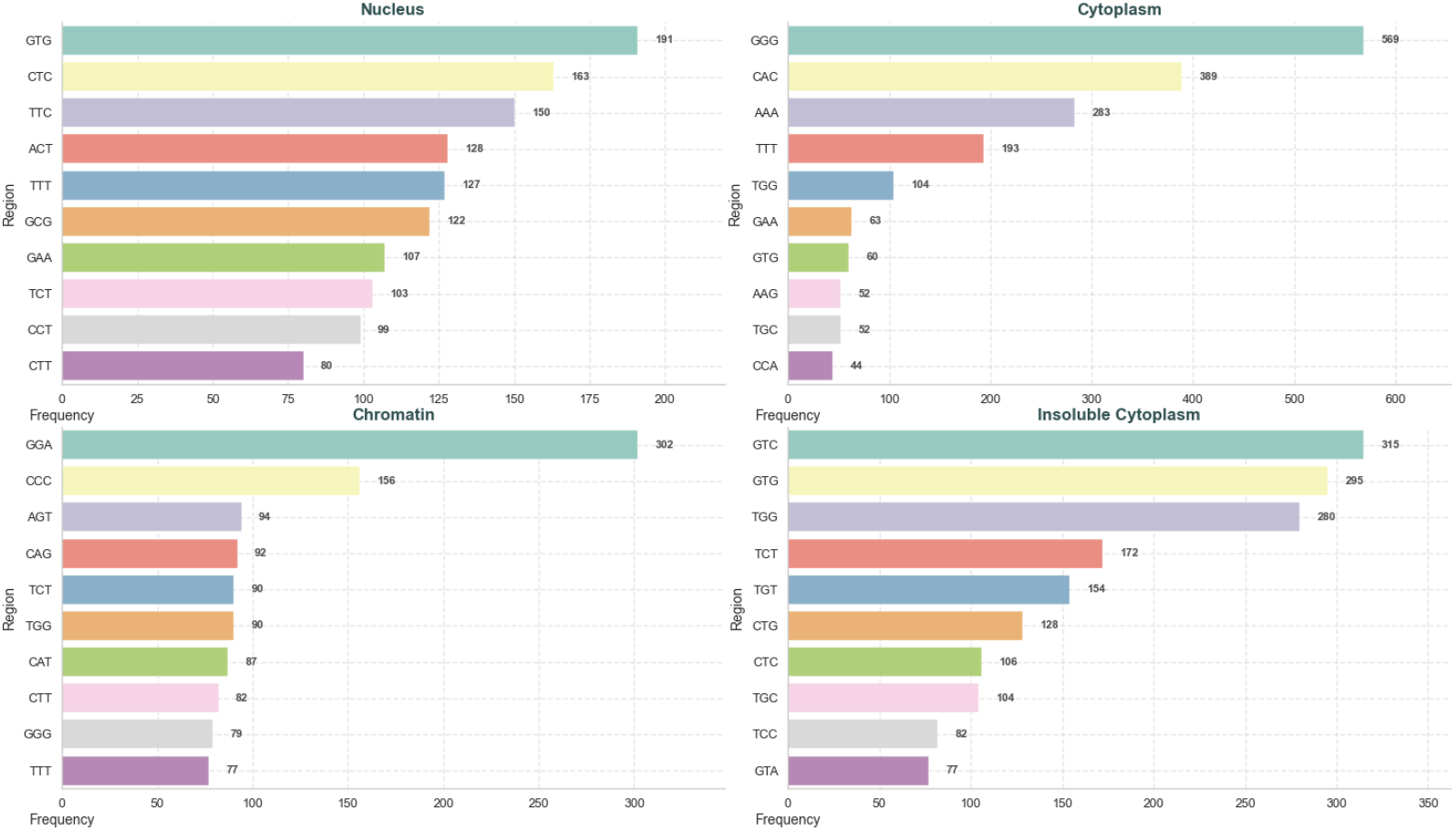
Top 10 motifs with the highest attention distributions in the four subcellular compartments

#### Sequence embedding

The paper adopts a subsequence embedding method [22]. The core idea is to divide the lncRNA sequence into a series of consecutive, non-overlapping subsequences, then extract patterns from each subsequence and combine these patterns to represent the complete lncRNA sequence. The specific steps are as follows:

- Split an lncRNA sequence into *n* consecutive and non-overlapping subsequences;
- Use the *word2vec* technique [19] to pre-train all lncRNA sequences in the dataset, extracting distributed representations of *k*-mers to represent subsequence features;
- Select an appropriate *k* value;
- Finally, combine these subsequence vectors to represent the complete lncRNA sequence.

#### Two-Layer FPN Network for Feature Extraction

After the embedding layer, we use a two-layer FPN network for feature extraction, employing convolutional kernels with different output channel numbers to efficiently capture multi-scale features [13]. The first convolutional layer has an input channel of 512, an output channel of 128, and a kernel size of 3, primarily extracting short-range local features while reducing the number of parameters and computational cost through dimensionality reduction. The second convolutional layer has an input channel of 128, with the output channel further reduced to 64 and a kernel size of 5, aiming to capture longer-range feature patterns and enhance the global representation ability of the features. Finally, by concatenating the original features with the multi-scale features extracted through convolution, the model integrates both local and global information to provide a more comprehensive representation of the input sequence.

#### Mamba2 block

As an emerging and efficient sequence modeling tool, the Mamba model [8] provides strong support for handling long-sequence tasks. We attempt to introduce the Mamba model into the lncRNA localization prediction task, leveraging its efficient modeling capabilities for long sequences to provide more comprehensive feature representations for localization prediction. Specifically, we use the Mamba2 block[7] with the following parameter settings: `d state=128`, `d conv=4`, `expand=2`, and `headdim=256`. These parameters were chosen based on our trials to achieve a balanced and efficient sequence representation.

#### Improved Localization-specific Attention Module

Building upon the Localization-specific Attention Module introduced in LncLocFormer [25], we propose an enhanced version designed to better address the multi-label classification task of lncRNA subcellular localization. The key innovation in our approach is the integration of prior knowledge about sequence motifs that are essential for specific subcellular localizations, allowing us to more accurately identify and focus on important nucleotides within each subcellular region.

In our improved attention mechanism, we still consider four primary subcellular regions, and for each region, we train a corresponding attention matrix *a*_1_, *a*_2_, *a*_3_, *a*_4_. These matrices allow the model to focus on the most relevant features for each localization.

The attention scores for each region are calculated using the following formula:

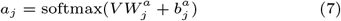

where *a*_*j*_ represents the attention score for region *j*, 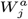 is the weight matrix for that region, and 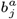 is the bias term.

Next, for each region, the corresponding attention score *a*_*j*_ is used to weight the feature matrix *V*, and the localization probability for that region is output through a fully connected layer and a *sigmoid* activation function:

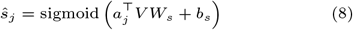

Finally, *ŝ*_*j*_ (*j* = 1, 2, 3, 4) represents the multi-label classification prediction for the four subcellular regions. The model parameters are updated by calculating the cross-entropy loss with the true labels.

To integrate existing research on lncRNA localization prediction and considering the importance of specific motifs in sequence function, we improved the deep pseudo-label attention mechanism by proposing a motif attention enhancement method. Specifically, we designed a function called *contains sequences* to detect whether the input sequence *tokenized seq* contains a predefined list of motifs.

The first motif we considered is the AGCCC motif, which acts as a general nuclear localization signal [27]. Additionally, for motifs of length 5, we referenced the AUUUA motif [3] as a potential motif that could have significant localization effects. However, after performing 5-mer encoding on the dataset, we found no occurrences of the selected motifs. Therefore, we switched to 3-mer encoding and, based on the results of LncLocator 2.0 [14], selected the top 10 nucleotide combinations that contributed the highest scores as candidate motifs. These nucleotide combinations are: TTT, AAA, TTG, TTA, GTT, ATT, TAA, GAA, AAT, and AAG. After performing 3-mer encoding, we found that the dataset contained 20 occurrences of the TTT motif, demonstrating the viability of this approach.

During the forward propagation of the model, if predefined motifs are detected in the input sequence, the attention weight for the region containing the motif is amplified by a factor of 1.5. Specifically, the attention weight for the motif region *α*_motif_ is updated according to the following formula:

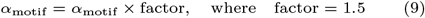

This mechanism dynamically adjusts the attention distribution, making the model focus more on key sequence patterns.

#### Loss Function

During training, we employed the basic cross-entropy loss function for optimization.

For each subcellular localization task, we use the *sigmoid* activation function to calculate the predicted probability *ŝ*_*j*_ for each region, and then compute the cross-entropy loss based on the difference between the predicted value and the true label:

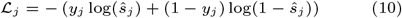

where *y*_*j*_ is the true label, and *ŝ*_*j*_ is the model’s predicted probability for the *j*-th subcellular region.

Since the lncRNA localization problem is a multi-label classification task, the total loss is the weighted sum of the losses for all subcellular regions:

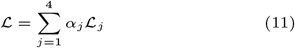

where *α*_*j*_ is the weight for each subcellular region, controlling the contribution of each region to the total loss. This loss function helps the model optimize the localization predictions for multiple subcellular regions simultaneously, enabling more accurate lncRNA localization.

## Result

### Evaluation Index

In this study, five evaluation metrics are used to assess the performance of the models: Average F1 (Ave-F1), Micro Precision (MiP), Micro Recall (MiR), Micro F1 (MiF), and Macro Area Under Curve (MaAUC). Each of these metrics provides a different perspective on the model’s performance in multi-label classification tasks.

**Average F1 (Ave-F1)** is the average of the F1 scores computed for each label. The F1 score is the harmonic mean of precision and recall and is calculated as:

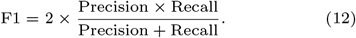

Ave-F1 represents the mean of the individual F1 scores across all labels.

**Micro Precision (MiP)** is computed globally by counting the total true positives (TP), false positives (FP), and false negatives (FN) across all classes. It is defined as:

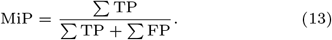

This metric reflects the model’s ability to predict positive labels across all classes.

**Micro Recall (MiR)** is calculated globally by considering the total true positives (TP) and false negatives (FN) across all classes. It is given by:

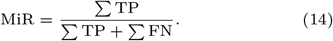

This metric assesses the model’s ability to identify all relevant instances.

**Micro F1 (MiF)** is the harmonic mean of Micro Precision and Micro Recall, providing a balanced evaluation of precision and recall at the global level. It is calculated as:

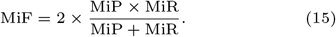

MiF combines the strengths of both precision and recall in a single metric.

**Micro Area Under Curve (MiAUC)** is the area under the Receiver Operating Characteristic (ROC) curve, calculated globally across all classes, and averaged by micro-averaging. Due to the data imbalance in Dataset 2, we use MiAUC instead of MaAUC to better reflect the model’s performance across the imbalanced data. In contrast, Dataset 1 employs the MaAUC metric to evaluate model performance, as it does not exhibit significant data imbalance.

**Macro Area Under Curve (MaAUC)** is the area under the Receiver Operating Characteristic (ROC) curve, averaged across all classes. A higher MaAUC indicates better model discrimination ability between positive and negative classes. The ROC curve plots the true positive rate (TPR) against the false positive rate (FPR) at various threshold settings.

### Comparison with Other Deep Learning Baseline Models

To evaluate the performance of LncMamba against other deep learning baseline models, we performed 5-fold cross-validation (5-fold CV). Specifically, the dataset was first split into a training set (90%) and a held-out test set (10%) for Dataset 1, while Dataset 2 was directly used as the training set. Then, the training set was further divided into 80% for training and 20% for validation, and 5-fold cross-validation was performed. This process was repeated five times, and the final prediction result was the average of the five validation outcomes. Additionally, the optimal MaAUC weight from the 5-fold cross-validation on Dataset 1 was applied to the test set results.

We compared some basic models with the latest popular models: LncLocFormer [25], DeepLncLoc [26], GraphLncLoc [11], CFPLncloc [18].For lncLocator 1.0 and SGCL-LncLoc, these are web-based models and therefore could not be trained on our dataset. As a result, we only performed comparisons on the test set.

The performance of LncMamba and the other baseline models using 5-fold CV is shown in Table 1 and Table 3. The results on the test set are shown in Table 2.The weights used for the related popular models are provided in Table 4.

**Table 1.** Model Performance Comparison on Dataset1.

| Model | Ave-F1 | MiP | MiR | MiF | MaAUC |
| --- | --- | --- | --- | --- | --- |
| K-mer + MLP[25] | 0.670 | 0.651 | 0.645 | 0.648 | 0.629 |
| Word2vec + MLP[25] | 0.690 | 0.664 | 0.660 | 0.662 | 0.582 |
| Word2vec + CNN + MLP[25] | 0.697 | 0.692 | 0.652 | 0.672 | 0.616 |
| Word2vec + Bi-LSTM + MLP[25] | 0.689 | 0.670 | 0.653 | 0.661 | 0.624 |
| GloVe + MLP[25] | 0.697 | 0.685 | 0.656 | 0.670 | 0.596 |
| GloVe + CNN + MLP[25] | 0.697 | 0.674 | 0.677 | 0.675 | 0.606 |
| GloVe + Bi-LSTM + MLP[25] | 0.679 | 0.712 | 0.602 | 0.652 | 0.582 |
| LncLocFormer[25] | 0.719 | 0.683 | 0.721 | 0.701 | 0.648 |
| DeepLncLoc[26] | 0.136 | 0.375 | 0.375 | 0.375 | 0.494 |
| GraphLncLoc [11] | 0.594 | 0.594 | 0.594 | 0.594 | 0.594 |
| CFPLncloc[18] | 0.651 | **0.750** | 0.531 | 0.319 | **0.694** |
| **LncMamba** | **0.732** | 0.653 | **0.791** | **0.712** | 0.562 |

**Table 2.** Model Performance Comparison on Test Set.

| Model | Ave-F1 | MiP | MiR | MiF | MaAUC |
| --- | --- | --- | --- | --- | --- |
| LncLocFormer[25] | 0.707 | 0.655 | 0.756 | 0.702 | 0.624 |
| DeepLncLoc[26] | 0.360 | 0.360 | 0.360 | 0.120 | **0.795** |
| GraphLncLoc [11] | 0.586 | 0.586 | 0.586 | 0.330 | 0.670 |
| lncLocator 1.0[30] | 0.290 | 0.500 | 0.500 | 0.500 | 0.550 |
| CFPLncloc[18] | 0.693 | **0.718** | 0.737 | 0.320 | 0.666 |
| **LncMamba** | **0.737** | 0.648 | **0.837** | **0.731** | 0.608 |

**Table 3.**
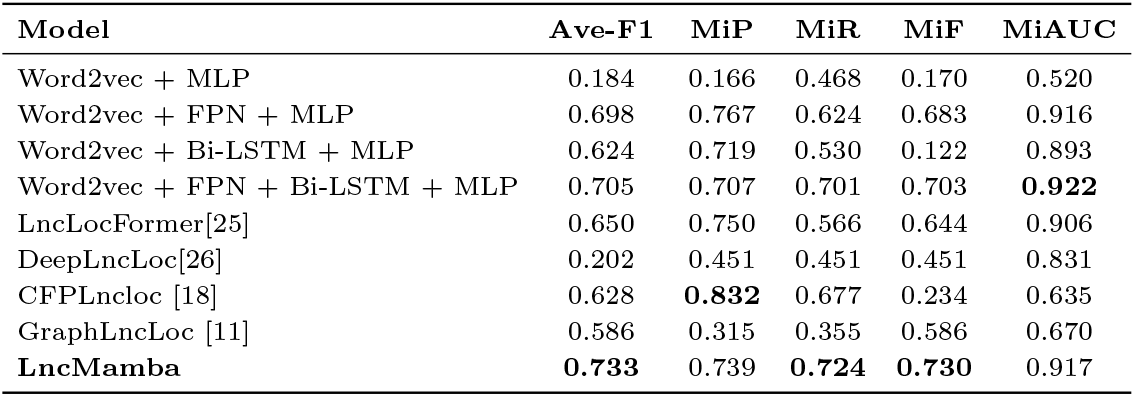
Model Performance Comparison on Dataset2.

**Table 4.** Model Weights Size Comparison.

| Model | Model Weights Size (MB) |
| --- | --- |
| LncLocFormer [25] | 25.46 |
| DeepLncLoc [26] | 0.33 |
| CFPLncloc[18] | 170.45 |
| GraphLncLoc [11] | 0.25 |
| **LncMamba** | **16.53** |

The 5-fold cross-validation results demonstrate that LncMamba outperforms other baseline models across most evaluation metrics on both datasets. Specifically, on Dataset 1, LncMamba achieves the highest Ave-F1 score of 0.732, surpassing models like LncLocFormer (0.719) and Word2vec + CNN + MLP (0.697). Additionally, LncMamba achieves the highest MiR score of 0.791 and MiF score of 0.712, showcasing its exceptional ability to accurately identify relevant features in the subcellular localization of lncRNAs.

On Dataset 2, LncMamba continues to excel, achieving an Ave-F1 score of 0.733, significantly outperforming models such as Word2vec + FPN + Bi-LSTM + MLP (0.705) and LncLocFormer (0.650). Moreover, it achieves the highest MiR score of 0.724 and MiF score of 0.730, demonstrating its strong performance on a dataset with longer sequences.

These advantages are maintained on independent test sets, where LncMamba achieves an average F1 score of 0.737, a MIP score of 0.731, and a MiR score of 0.837, the highest among all models.

Regarding model size, LncMamba demonstrates relatively efficient memory usage. It has a model size of 16.53 MB, significantly smaller than CFPLncloc (170.45 MB), making it a more computationally efficient option. It is also more compact compared to LncLocFormer (25.46 MB), while still maintaining competitive performance.

### Ablation Study

To evaluate the contributions of key components in the LncMamba model, we conducted two ablation studies. The first study involves removing the two-layer FPN network, and the second study examines the model’s performance without the improved attention mechanism.

Ablation of Two-Layer FPN Network: In this experiment, we removed the two-layer FPN network to assess the contribution of multi-scale feature extraction to the overall model performance.

Ablation of Improved Attention Mechanism: In the second experiment, we excluded the improved localization-specific attention mechanism, leaving the model to rely on the original attention mechanism.

The performance of LncMamba after these modifications was evaluated using the same five metrics as in the previous section: Average F1 (Ave-F1), Micro Precision (MiP), Micro Recall (MiR), Micro F1 (MiF), and Macro Area Under Curve (MaAUC). The results of the ablation experiments are summarized in Table 5.

**Table 5.** Ablation Study Results.

| Model | Ave-F1 | MiP | MiR | MiF | MaAUC |
| --- | --- | --- | --- | --- | --- |
| LncMamba (Full Model) | **0.732** | 0.653 | **0.791** | **0.712** | 0.562 |
| No FPN (Ablation 1) | 0.726 | 0.658 | 0.765 | 0.706 | 0.573 |
| No Attention (Ablation 2) | 0.725 | **0.668** | 0.741 | 0.701 | **0.580** |

Removing the FPN network (Ablation 1) caused a slight drop in all metrics, particularly in Ave-F1 (0.732 to 0.726). This suggests that multi-scale feature extraction via the FPN network contributes to small but significant improvements in performance, especially in MiP and MiR.

Excluding the improved attention mechanism (Ablation 2) resulted in a minor decrease across most metrics, with Ave-F1 remaining stable at 0.725. However, MiP and MiR dropped slightly, while MaAUC improved to 0.580. This suggests that the attention mechanism enhances the model’s sensitivity to specific motifs, but its removal does not drastically affect the model’s overall performance.

Overall, the full model consistently outperforms the ablation variants, highlighting the complementary contributions of the two-layer FPN network and the attention mechanism for optimal performance in lncRNA subcellular localization prediction.

### Exploration of Localization Motifs

To explore the motifs that play a crucial role in lncRNA subcellular localization, we conducted a statistical analysis of the attention distribution across four subcellular compartments: nucleus, cytoplasm, chromatin, and insoluble cytoplasm. Specifically, we focused on the 3-mer sequences with the highest attention scores and their surrounding sequences, identifying the top motifs in each compartment based on the frequency of occurrence.

The model was trained using the unmodified deep pseudo-label attention mechanism, and the model weights with the highest MaAUC from the 5-fold cross-validation were selected for this analysis. The resulting motif distributions were derived from a comprehensive statistical analysis performed on a dataset containing 811 lncRNAs.

From the analysis, we observed several key motifs in different subcellular compartments. In the nucleus, GTG, CTC and TTC motifs appeared the most, 191 times, 163 times and 150 times, respectively, indicating that these sequences are important for nuclear localization. The high frequency of GTG and CTC motifs may indicate that they are involved in interactions with nuclear proteins or structures. In the cytoplasm, GGG and CAC moths appeared 569 times and 389 times, respectively. The widespread existence of G - and C-rich moths suggests that they are involved in RNA stability and translation regulation, which is essential for the normal function of lncRNA in the cytoplasm. In the chromatin compartment, motif GGA appeared the most frequently (302 times), which may be involved in RNA-protein interactions that regulate chromatin structure or gene expression. In the insoluble cytoplasm, GTC, GTG, TGG and other motifs are highly representative and appear more frequently than other motifs. These motifs may be related to the stability of the insoluble part of the cytoplasm and specific protein interactions.

Overall, different motifs were identified in each cell compartment, GTG and CTC motifs were enriched in the nucleus, GGG and AAA dominated in the cytoplasm, GGA was favored in the chromatin compartment, and GTC and GTG motifs were enriched in the insoluble cytoplasm. These findings highlight the important role of specific moieties in lncRNA subcellular localization and provide new insights into their functional mechanisms.

## Conclusion

In this study, we addressed the problem of lncRNA subcellular localization prediction by proposing a novel deep neural network based on the Mamba model. We conducted an in-depth exploration of feature extraction methods, drawing inspiration from the concept of the FPN network. We designed a two-layer FPN network to extract multi-scale features, effectively capturing both local and global information. For the backbone network, we applied the Mamba network to the task of lncRNA subcellular localization prediction for the first time, achieving competitive performance.

Furthermore, we incorporated prior knowledge to improve the localization-specific attention mechanism. This enhancement allowed the model to focus more on key motifs, thereby improving classification accuracy and enabling better model interpretability.

In the exploration of localization motifs, we performed a statistical analysis of the motifs across different subcellular compartments. The results revealed distinct patterns: the nucleus is enriched with motifs containing T, C, and G; the cytoplasm is dominated by motifs with G and A; in the insoluble cytoplasm, motifs containing G and T are prevalent; and in chromatin, motifs with C and G are enriched. These findings underscore the significant connection between nucleotide motifs and lncRNA subcellular localization. They provide valuable insights into the functional roles of specific motifs, opening new avenues for further research in understanding the mechanisms behind lncRNA localization and its biological implications.

## References

1. M. N. Asim, M. A. Ibrahim, M. I. Malik, C. Zehe, and A. Dengel. El-rmlocnet: An explainable lstm network for rna-associated multi-compartment localization prediction. Computational and Structural Biotechnology Journal, 20:1737–1750, 2022.

2. T. Bai and B. Liu. ncrnalocate-el: a multi-label ncrna subcellular locality prediction model based on ensemble learning. Briefings in Functional Genomics, 22(5):442–452, 2023.

3. Christine Barreau, Laurent Paillard, and Howard B. Osborne. Au-rich elements and associated factors: are there unifying principles? Nucleic Acids Research, 33(22):7138–7150, 2006.

4. Mary Catherine Bridges, Amanda C. Daulagala, and Antonis Kourtidis. Lnccation: lncrna localization and function. Journal of Cell Biology, 220(2):e202009045, 01 2021.

5. Robert S. Bucy and Brian D. O. Anderson. Discrete-Time Control Systems. Addison-Wesley, 1968.

6. Claudia Carrieri, Laura Cimatti, Marta Biagioli, Anne Beugnet, Silvia Zucchelli, Stefania Fedele, Elisa Pesce, Isidre Ferrer, Licio Collavin, Claudio Santoro, Alistair R. R. Forrest, Piero Carninci, Stefano Biffo, Elia Stupka, and Stefano Gustincich. Long non-coding antisense rna controls uchl1 translation through an embedded sineb2 repeat. Nature, 491(7424):454–457, 2012.

7. Tri Dao and Albert Gu. Transformers are ssms: Generalized models and efficient algorithms through structured state space duality. arXiv preprint arXiv:2405.21060, 2024.

8. A. Gu and T. Dao. Mamba: Linear-time sequence modeling with selective state spaces. arXiv preprint arXiv:2312.00752, 2023.

9. Rudolf E. Kalman. A new approach to linear filtering and prediction problems. Transactions of the ASME–Journal of Basic Engineering, 82(1):35–45, 1960.

10. Min Li, Baoying Zhao, Yiming Li, Pingjian Ding, Rui Yin, Shichao Kan, and Min Zeng. Sgcl-lncloc: An interpretable deep learning model for improving incrna subcellular localization prediction with supervised graph contrastive learning. Big Data Mining and Analytics, 7(3):765–780, 2024.

11. Min Li, Baoying Zhao, Rui Yin, Chengqian Lu, Fei Guo, and Min Zeng. Graphlncloc: long non-coding rna subcellular localization prediction using graph convolutional networks based on sequence to graph transformation. Briefings in Bioinformatics, 24(1), January 2023.

12. T. Y. Lin, P. Dollár, R. Girshick, K. He, B. Hariharan, and S. Belongie. Feature pyramid networks for object detection. In Proceedings of the IEEE conference on computer vision and pattern recognition, pages 2117–2125, 2017.

13. Tsung-Yi Lin, Piotr Dollár, Ross Girshick, Kaiming He, Bharath Hariharan, and Serge Belongie. Feature pyramid networks for object detection. In Proceedings of the IEEE Conference on Computer Vision and Pattern Recognition (CVPR), pages 2117–2125, 2017.

14. Yang Lin, Xiaoyong Pan, and Hong-Bin Shen. lnclocator 2.0: a cell-line-specific subcellular localization predictor for long non-coding rnas with interpretable deep learning. Bioinformatics, 37(16):2308–2314, 2021.

15. F. Liu, L. Zhang, J. Chen, X. Wang, X. Wang, and S. Li. Locate-r: subcellular localization of long non-coding rnas using nucleotide compositions. Genomics, 112(3):2583–2589, 2020.

16. Lennart Ljung. System identification: Theory for the user. Prentice Hall, 1999.

17. Jeffrey R. Moffitt and Xiaowei Zhuang. Rna imaging with multiplexed error-robust fluorescence in situ hybridization (merfish). Methods in Enzymology, 572:1–49, 2016.

18. Sheng Wang, Zu-Guo Yu, Guo-Sheng Han, and Xin-Gen Sun. Cfplncloc: A multi-label lncrna subcellular localization prediction based on chaos game representation and centralized feature pyramid. International Journal of Biological Macromolecules, 297:139519, 2025.

19. Y. Wu, M. Gao, M. Zeng, et al. Bridgedpi: a novel graph neural network for predicting drug-protein interactions. Bioinformatics, 38(10):2571–2578, 2022.

20. Y. Xu, X. Zhao, S. Liu, and W. Zhang. Predicting long non-coding rnas through feature ensemble learning. BMC Genomics, 2020.

21. Guodong Yang, Xiaozhao Lu, and Lijun Yuan. Lncrna: A link between rna and cancer. Biochimica et Biophysica Acta (BBA) - Gene Regulatory Mechanisms, 1839(11):1097–1109, 2014.

22. M. Zeng, C. Lu, F. Zhang, et al. Sdlda: lncrna-disease association prediction based on singular value decomposition and deep learning. Methods, 179:73–80, 2020.

23. M. Zeng, Y. Wu, Y. Li, R. Yin, C. Lu, J. Duan, and M. Li. Lnclocformer: a transformer-based deep learning model for multi-label lncrna subcellular localization prediction by using localization-specific attention mechanism. Bioinformatics (Oxford, England), 39(12):btad752, 2023.

24. Meng Zeng, Chao Lu, Zhi Fei, et al. Dmflda: a deep learning framework for predicting lncrna-disease associations. IEEE/ACM Transactions on Computational Biology and Bioinformatics, 18:2353–2363, 2021.

25. Min Zeng, Yifan Wu, Yiming Li, Rui Yin, Chengqian Lu, Junwen Duan, and Min Li. Lnclocformer: a transformer-based deep learning model for multi-label lncrna subcellular localization prediction by using localization-specific attention mechanism. Bioinformatics, 39(12):btad752, December 2023.

26. Min Zeng, Yifan Wu, Chengqian Lu, Fuhao Zhang, Fang-Xiang Wu, and Min Li. Deeplncloc: a deep learning framework for long non-coding rna subcellular localization prediction based on subsequence embedding. Briefings in Bioinformatics, 23(1):bbab360, 2022.

27. Bo Zhang, Lalith Gunawardane, Farzad Niazi, et al. A novel rna motif mediates the strict nuclear localization of a long noncoding rna. Molecular and Cellular Biology, 34:2318–2329, 2014.

28. Tao Zhang, Peng Tan, Lihua Wang, Nian Jin, Yong Li, Lining Zhang, and Zhi Hu. Rnalocate v2.0: an updated resource for rna subcellular localization with increased coverage and annotation. Nucleic Acids Research, 49(D1):D1065–D1071, 2021.

29. Y. Zhang, X. Li, X. Li, D. Wang, L. Zhang, and M. Zhang. Lnclocpred: predicting lncrna subcellular localization using multiple sequence feature information. IEEE Access, 8:124702–124711, 2020.

30. Cao Zhen, Xiaoyong Pan, Yang Yang, Yan Huang, and Hong-Bin Shen. The lnclocator: a subcellular localization predictor for long non-coding rnas based on a stacked ensemble classifier. Bioinformatics (Oxford, England), 34, 02 2018.

31. L. Zhu, H. Chen, and S. Yang. Lncsl: A novel stacked ensemble computing tool for subcellular localization of lncrna. International Journal of Molecular Sciences, 25(24), 2024.

